# One of the gang: social group dynamics in a juvenile passerine bird

**DOI:** 10.1101/456376

**Authors:** Victoria R. Franks, John G. Ewen, Mhairi McCready, J. Marcus Rowcliffe, Donal Smith, Rose Thorogood

**Affiliations:** Department of Zoology, University of Cambridge, Downing Street, Cambridge, CB2 3EJ, U.K.; Institute of Zoology, Zoological Society of London, Regent’s Park, London, NW1 4RY, U.K.; School of Environment & Life Sciences, University of Salford, Salford, M5 4WT, U.K.; Helsinki Institute of Life Science (HiLIFE), University of Helsinki, Finland; Research program in Organismal and Evolutionary Biology, Faculty of Biological and Environmental Sciences, University of Helsinki, Finland.

## Abstract

Living in groups comes with many potential benefits, especially for juveniles. Naïve individuals may learn how to forage, or avoid predators through group vigilance. Understanding these benefits, however, requires an appreciation of the opportunities juveniles have to associate with (and learn from) others. Here we describe social groups in terms of residency, movement, relatedness, and social associations from the perspective of juvenile hihi, a threatened New Zealand passerine bird. Over three years, we identified individuals in groups, their relatedness, and behavioural interactions. Using multistate analysis, we compared movement and residency of adults and juveniles and found that groups were composed predominately of juveniles which remained at group sites for longer than more transient adults. Movement of juveniles between groups did occur but was generally low. There was no evidence that siblings and parents were likely to be seen in groups together. With an initial understanding of group structure, we next asked what characteristics predicted assortment in social network associations. By identifying groups of co-occurring juveniles from time-stamped observations of individual hihi and building a social network, we found that juveniles were most likely to associate with other juveniles. Associations were also predominantly based on locations where hihi spent the most time, reflecting limited movement among separate groups. We suggest groups are best described as “gangs” where young hihi have little interaction with adults. These spatially-separated groups of juveniles may have consequences for social information use during the first few months of independence in young birds.

## Introduction

Social groups are found across the animal kingdom (Ward and Webster, 2016). Generally, group living is thought to provide advantages such as better foraging opportunities if groups can collectively out-compete other more dominant animals, and protection from predators through shared vigilance (Rubenstein, 1978; Molvar and Bowyer, 1994; Gompper, 1996; Hass and Valenzuela, 2002; Le Bohec, Gauthier-Clerc and Le Maho, 2005). Sociality may benefit juveniles in particular, as they are inexperienced compared to adults (Galef and Laland, 2005) and can be less adept at finding food (Sullivan, 1989; Lind and Welsh, 1994; Franks and Thorogood, 2018) or avoiding predators by themselves (Naef-Daenzer, Widmer and Nuber, 2001). Therefore, juveniles may be able to use information from others in groups (“social information”) to reduce their uncertainty about how best to behave (Dall *et al.*, 2005). However, who juveniles encounter in groups will affect the opportunities they have to learn socially (Seppänen *et al.*, 2007; Krause *et al.*, 2015), along with additionally impacting on other consequences of group living such as risk of contracting disease (Godfrey *et al.*, 2009; Drewe, 2010). Therefore, the dynamics of groups (where groups form, when they form, and which individuals group) need to be quantified to understand why animals are social, especially juveniles (Krause and Ruxton, 2002; Ward and Webster, 2016).

In young wild birds, there are examples of three types of groups which vary in age structure, relatedness structure, and site stability. Firstly, birds are well-known to form mobile foraging units or “flocks”, with no particular structure by relatedness or age (so are not unique to juveniles) (Morse, 1978; Saitou, 1978, 1979; Ekman, 1989; Templeton *et al.*, 2012). All group members can access ephemeral food sources as the group moves across an environment, and they also share costs of predator vigilance (Rubenstein, 1978; Molvar and Bowyer, 1994; Hass and Valenzuela, 2002; Sutton, Hoskins and Arnould, 2015). Secondly, juveniles can form groups without adults, such as “gangs” in ravens (*Corvus corax*) (Dall and Wright, 2009). Gangs are similar to flocks in that their main function is to access ephemeral food resources; however, gangs operate around stable roosting sites which act as information centres (Dall and Wright, 2009) and also allow juveniles to out-compete more dominant adults (Wright, Stone and Brown, 2003; Ward and Zahavi, 2008; Dall and Wright, 2009). Thirdly, juveniles may form stable congregations as “crèches”. These groups contrast with gangs or flocks because crèches form before juveniles become fully independent from parents, and serve to promote juvenile survival as parents actively care for their young with food provisioning and vigilance (Balda and Balda, 1978; Marzluff and Balda, 1992; Clayton and Emery, 2007). Despite group structure being a crucial part of the social environment of juveniles, little is known outside of these examples. Overall, juvenile behaviour is understudied in general, especially in birds (Templeton *et al.*, 2012).

One way groups may help juveniles overcome their naivety is by providing opportunities to learn rapidly when interacting with group members (Coussi-Korbel and Fragaszy, 1995; Krause and Ruxton, 2002). If groups are comprised of different age classes, or individuals that use the environment in different ways, juveniles may encounter a range of potential social information (Seppänen *et al.*, 2007; Pinter-Wollman *et al.*, 2013). For example, juveniles in flocks encounter both experienced adults and other juveniles, so information about ephemeral food sources can be shared among group members (“oblique” and “horizontal” transmission, respectively (van Schaik, 2010)). In gangs where associations are largely between juveniles only, interactions can include “social play” (documented in species such as ravens (Heinrich and Smolker, 1998)), where at least two individuals engage in a reciprocated behaviour and alternate between roles, and potentially share information (Diamond and Bond, 2003). However, the presence of many naïve individuals in gangs could increase the risk of associating with misinformed peers, especially if some individuals are more social than others (Pruitt *et al.*, 2016). Across the animal kingdom, genetically-related groups such as crèches promote associations between parents and offspring (Balda and Balda, 1978; Clayton and Emery, 2007) that allow for learning (e.g. European shags *Phalacrocorax aristotelis*: Velando, 2001; ravens *Corvus corax:* Schwab *et al.*, 2008; vervet monkeys *Chlorocebus pygerythrus:* van de Waal, Bshary and Whiten, 2014) and can even facilitate teaching (e.g. meerkats *Suricata suricatta* (Thornton, 2006; Thornton and Raihani, 2010)). Alternately, some studies suggest associations with non-kin can still be beneficial as these individuals may have a different range of experiences (Hatch and Lefebvre, 1997). Describing group structures and understanding how these affect associations should therefore help us to understand the benefits of group living for juveniles more clearly (Sih, Hanser and McHugh, 2009).

Analysing group behaviour in space and time can quantify broad-scale consistencies to show who groups, when and where. However, it does not fully capture how individuals interact as a consequence of group structure. Social network analysis can overcome this problem (Wey *et al.*, 2008); animals form social networks through non-random preferred and avoided associations which can be quantified and analysed statistically (Krause and Ruxton, 2002; Krause, Lusseau and James, 2009; Sih, Hanser and McHugh, 2009; Krause *et al.*, 2015). Therefore, here we first used a form of re-sighting analysis to consider movement, residency and relatedness in groups of juvenile and adult birds. We then compiled social networks to investigate how movement, residency and relatedness affected associations between individuals. Finally, we observed interactions between grouping individuals to understand how social behaviour may influence information sharing. Our study species was the hihi (*Notiomystis cincta*), an endemic New Zealand passerine. Hihi provide a good example where juveniles are known anecdotally to form groups during early life, although these have not been studied systematically before. We aimed, therefore, to describe group formation and membership, compare them to juvenile groups in other species, and understand how group characteristics affected associations (Table 1). If hihi groups were crèches, we predicted both adults (parents) and juveniles (siblings) to be consistently sighted together in groups. However, we would expect different structure if groups were gangs (juveniles should be present much more than adults) or flocks (individuals would not remain in one site; adults and juveniles would be present but unrelated).

**Table 1.** Predictions for group structure and social associations for juvenile hihi, with reference to previously-described groups of birds. All group types can be compared to a “null” unstructured group of randomly-associating individuals.

| Group characteristics | Group type |  |  |  |
| --- | --- | --- | --- | --- |
|  | Flock<br>(Saitou<br>1978;<br>1979) | Gang<br>(Marzluff<br>et al.,<br>1996) | Crèche<br>(Marzluff<br>& Balda<br>1992) | Null |
| (1) Age composition: |  |  |  |  |
| Juveniles are more resident in groups than adults |  |  |  |  |
| • Juveniles re-sighted more days than adults; | No | Yes | No | No |
| • Age structures associations |  |  |  |  |
| • Juveniles interact |  |  |  |  |
| (2) Spatial structure: |  |  |  |  |
| Juveniles group in consistent locations |  |  |  |  |
| • Low movement between separate groups; | No | Yes | Yes | No |
| • Location structures associations |  |  |  |  |
| (3) Relatedness: |  |  |  |  |
| Groups contain parents and offspring |  |  |  |  |
| • Juveniles consistently sighted with<br>parents/siblings; | No | No | Yes | No |
| • Relatedness structures associations |  |  |  |  |
| (4) Groups random (across/within years) | No | No | No | Yes |

## Methods

### STUDY POPULATION

Our study was conducted over three years (2015 – 2017) on Tiritiri Matangi Island (Auckland, New Zealand, 36°36'00.7"S 174°53'21.7"E), between January – April when juvenile hihi (birds in their first year) had fledged and dispersed from nests. This 2.5km^2^ island is characterised by a central longitudinal ridge (60-80m altitude) with a series of latitudinal ridges and gullies on either side covered in a mixture of original and replanted native bush. Supplementary sugar water feeders are provided year-round for hihi at five sites across the island. This is a closed population with no immigration or emigration (except through birth and mortality) and all individuals are uniquely identifiable from coloured leg ring combinations. The population varied between 180 and 270 individuals over the three years, with similar proportions of juveniles and adults (second year or older) each year (Smith and Ewen, 2015; McCready and Ewen, 2016, 2017). Every year, all breeding attempts are monitored and identities of breeding pairs recorded. During this study, parentage and siblings were assigned based on social relationships. Although there is variable extra-pair paternity in hihi (Ewen, Armstrong and Lambert, 1999; Brekke *et al.*, 2013), all nest-mates were most likely to be at least maternal-siblings (there is no evidence of conspecific brood parasitism in hihi) and the social male cares for the offspring in his nest (Ewen and Armstrong, 2000). The first year of our study (2015) was a poorer breeding season than 2016 and 2017 (2015: 89 fledglings; 2016: 132 fledglings; 2017: 151 fledglings); thus, we accounted for year in any analyses using combined data.

### DETECTING GROUPS

Each year, we surveyed for groups between January – February in spatially-separated areas of forest habitat. In 2016 and 2017, we increased the search area to ensure no other potential groups were missed. The numbers of unique juveniles were recorded for one hour in each location, and group sites were assigned after two weeks if we saw at least three juveniles during more than 80% of 10 surveys per location. We further confirmed that there were no other sites with higher numbers of juveniles during the annual February census of the population, which is conducted every year by trained conservation staff who survey the entire island over 40 hours. We then continued to survey group locations from February – April, using one-hour surveys divided into 30-second time blocks (one survey = 120 blocks). Within each block we recorded the identity of all hihi (both juvenile and adult) perched within a 10-metre radius of the observer. We recorded individuals present across blocks to determine presence to the nearest 30 seconds, and also the occurrence of behavioural interactions and the identities of the individuals involved (Table 2; Figure 1). Interactions were classed as “directed” if there were clear initiators; however, for some behaviours individuals were only ever seen to interact equally, so we classified these as “undirected” (Table 2). All observations were made with binoculars (Zeiss Conquest^®^ HD 8×42) by one observer (VF). In total we recorded 15 hours per group site in 2015, and 25 hours per site in 2016 and 2017; surveys were distributed evenly across the three months.

**Figure 1.**
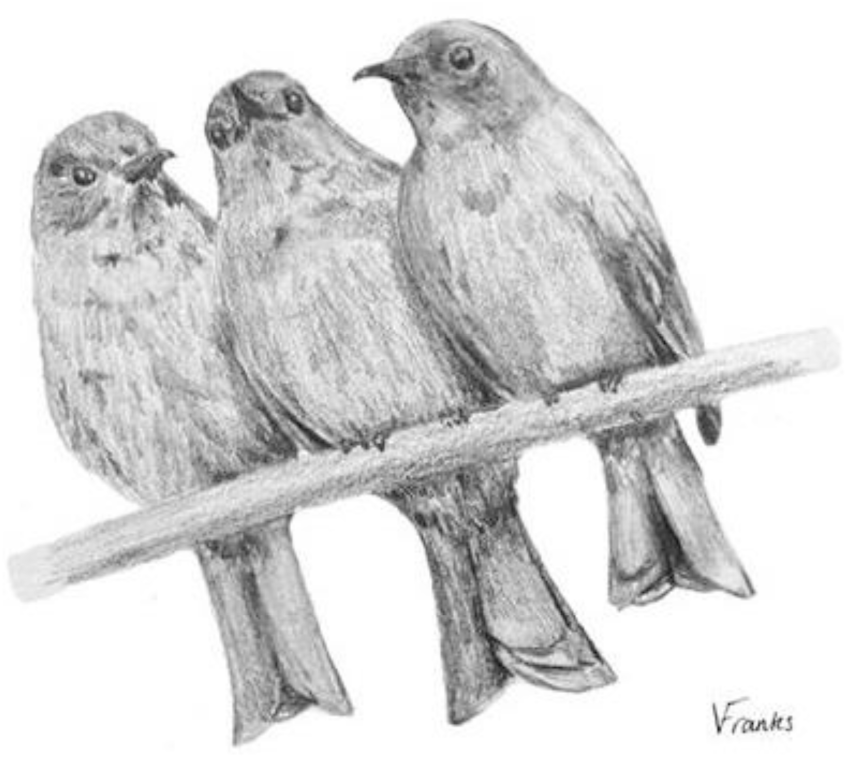
Sketch of huddle behaviour, with three juvenile hihi perching side-by-side on a branch.

**Table 2.** Ethogram of common interactions observed between juveniles in groups, and their definitions.

|  | Interaction | Description |
| --- | --- | --- |
| Directed behaviours | Chase | Individual moves towards another perched hihi, displaces it, and then continues moving in the same direction as the second bird. |
|  | Chased | Individual moves away from its original perch after another hihi has initiated moving towards the focal individual |
|  | Follow | Individual leaves a perch to move in same direction as another hihi that moves off before focal individual. |
|  | Followed | Individual leaves a perch before another perched bird, and the second bird moves off in the same direction as the focal individual. |
| Undirected behaviours | Huddle | Two or more birds perch touching side-by-side and do not move from position on perch. May include allopreening. |
|  | Playfight | Two or more birds perch touching side-by-side, then peck at each other, hang up-side down on branch, shuffle next to each other along branch. |

### DATA ANALYSIS

#### Re-sighting and movement analysis

We first used a multistate approach in Program MARK (version 9.0) (White and Burnham, 1999) to understand residency in group sites and movement between sites for juveniles and adults. Multistate analyses estimate survival (*S*) and re-sighting (*ρ*) of identifiable individuals of different “states” from repeated sightings across replicated surveys, along with likelihood of movement between states (*ψ*) (White, Kendall and Barker, 2006). Survival and re-sighting are often inherently linked and considered together to investigate population dynamics (i.e. to identify true mortality instead of absence of detection). However, in our study we used these three parameters in a novel way to determine different patterns of hihi group structure. We assumed mortality was constant over our short study periods each year (Jolly, 1982), supported by previous studies of adult and juvenile survival in this population (Low and Pärt, 2009). Including varying survival by age in preliminary exploratory models also indicated low mortality (98-99% survival each year for both ages across all observational surveys; results not presented) suggesting birds were alive throughout the study period. We could then use *ρ* to quantify presence/absence in groups (larger values of *ρ* indicated high residence within groups as opposed to independent living), and track movement between groups using state transitions (higher values of *ψ* indicating greater movement between known groups). In 2016 we verified that our survey method was reliable (see Appendix) to be confident in estimating *ρ*.

We constructed re-sighting histories for each bird seen each year to represent if, and in which group, it was seen. Different groups were not surveyed at the same time (due to one observer), so we combined surveys together to create occasions that represented every group site. There was a maximum of two days between combined surveys to limit movement between groups within a survey occasion; if this did occur we took the newest site as the site of residence for that individual in that occasion to account for movement (this occurred rarely: 2015 = 1/656 re-sightings; 2016 = 9/1974 re-sightings; 2017 = 7/3180 re-sightings). Thus, there were 8 survey occasions in 2015, 14 in 2016, and 20 in 2017. An example re-sighting history for one individual in 2015 is “aa0abbbb”, where the bird was seen in group “a” in survey occasions 1 and 2, not seen in survey occasion 3, seen in group “a” in survey occasion 4, and seen in group “b” for the remainder of survey occasions. We also specified if an individual was juvenile or adult with its re-sighting history. Therefore, our general starting model using our encounter histories in all three years was:

#### *S*(.) *ρ*(age*survey occasion) *ψ*(age + group)

Here, we quantified different residency in groups between adults and juveniles by assessing if re-sighting *ρ* varied across each survey occasion and between age groups, and if movement *ψ* between groups also varied with age. For *ψ*, we specified group differences as varying distance and topology between groups could affect likelihood of moving between each group (Martin *et al.*, 2006; Strandburg-Peshkin *et al.*, 2017). Finally, different intervals between survey occasions were accounted for so likelihoods were not confounded by time.

Assessing fit allows for accurate inference from more reduced models (Burnham and Anderson, 2002; Cam *et al.*, 2004; Martin *et al.*, 2006). Therefore we assessed goodness-of-fit (GOF) of our starting models each year using median ĉ (variance inflation factor, a measure of overdispersion), which is generated by assessing the distribution of model deviances (White and Burnham, 1999; Gath, 2017). Values of median ĉ > 1 suggest overdispersion that needs to be corrected for in analyses while values > 4 suggest a structural failure of the general model (Burnham and Anderson, 2002). Each year, there were low levels of overdispersion (median ĉ: 2015 = 1.51; 2016 = 1.39; 2017 = 1.72) so we used the value of median ĉ as a correction factor for all further multistate analyses.

For each year, we constructed sets of models with all possible combinations of *ρ* and *ψ* parameters and ranked these by their corrected Quasi-Akaike Information Criterion (QAICc) values. AICc values represent the change in fit in comparison to the top-ranked model, and QAICc is used when ĉ is corrected following GOF testing. Any model < 2 QAICc units from the top-ranked model were considered equally well-supported. We also calculated QAICc weights for each model based on change in QAICc value from top-ranked model, which gave the relative likelihood that it was the most appropriate model (Burnham and Anderson, 2002). Any parameters included in models with QAICc weight > 0.00 were included in model-averaging to calculate effect sizes and 95% confidence intervals. Any parameter with a confidence interval that did not include 0.00 was considered to have a significant effect.

All further analyses were conducted in R (version 3.5.0) (R Core Team, 2017). To determine if juveniles were using the same group sites as their parents and maternal-siblings, using each juvenile’s re-sighting history we calculated, per bird, the proportion of occasions it was seen in the same group as either of its parents, and the proportion of occasions it was seen in the same group as its maternal-siblings. We excluded any juveniles seen in one survey only as we could not calculate a proportion for these (*N*: 2015 = 10; 2016 = 18; 2017 = 10). When calculating proportions of time spent with maternal-siblings we also excluded any juveniles from single-fledgling nests or those with no maternal-siblings seen during our surveys (which may have died after fledging) (*N*: 2015 = 7; 2016 = 10; 2017 = 8). We assessed if juveniles that grouped closer to their nest-of-origin were more likely to co-occur with their maternal-siblings. We used a binomial Generalised Linear Model (GLM) where proportion of surveys with maternal-siblings was the response variable; using a proportion meant we could analyse all years together. Our predictors were proximity to nest-of-origin (distance to nearest 50m from group site to nest-of-origin, measured using Google Maps), number of surveys to ensure co-occurrence with maternal-siblings was not due to sampling bias, and year of survey (2015, 2016, 2017) to compare patterns among years. We constructed a set of candidate models including all combinations of predictors and ranked models by their AICc values. For any model < 2 AICc units larger than the top-ranked model, we calculated averaged effect sizes (±95% confidence intervals) for predictors using the package AICcmodavg (version 2.1-1) (Mazerolle, 2017). Based on the evidence from this initial exploration (see Results) we did not analyse effects of relatedness further using MARK, or in social network analysis.

#### Social network analysis

We constructed a social network for each year separately using the R package asnipe (version 1.1.9) (Farine, 2013). First we used the “gmmevents” function to detect temporal clusters in our time-stamped (to within 30s) sightings data and build an association matrix (Farine, 2013; Psorakis *et al.*, 2015). Using this approach avoids artificially restricted associations, which can occur using a more fixed time-window approach (Psorakis *et al.*, 2015). To validate if “gmmevents” groups represented true associations, we then compared the length of time (number of sequential observation blocks) we re-sighted hihi during observations to event lengths generated by “gmmevents”. All networks were weighted, which incorporates both the number and strength of social connections and are considered more robust than binary networks (Farine, 2014). Any hihi with fewer than 3 observation records were not included in networks (juvenile *N*: 2015 = 6; 2016 = 1; 2017 = 4; adult *N*: 2015 = 12; 2016 = 8 2017 = 7).

As network data is not independent and thus violates the assumptions of many statistical tests, we compared observed networks to randomised networks as a null model to test hypotheses (Croft *et al.*, 2011; Farine and Whitehead, 2015; Farine, 2017). All randomised networks were generated using permutations of the data-stream in asnipe, which randomly swaps records of individuals and is considered best practice instead of node-based permutations because it maintains original data structure and controls for sampling bias (Farine, 2013, 2014, 2017; Farine and Whitehead, 2015). Significance was calculated by dividing the number of times the test statistic of the real network was smaller than the test statistics of randomised networks by 1000 (the number of permutations). All *P-*values generated using random networks comparisons are specified here as *P*_rand_. Visualisations of networks were constructed in Gephi (version 0.9.2) (Bastian, Heymann and Jacomy, 2009) with a force-atlas layout that clustered together more strongly associating nodes.

We tested if hihi formed non-random associations in their groups compared to permuted networks using the coefficient of variation (“cv”). The value of cv describes variation in edge weights across a network: extreme values of cv are 0 and 10, but any values over 0.6 are considered to represent differentiated networks (groups are comprised of strong, repeated connections) (Farine and Whitehead 2015). We then explored if non-random associations were explained by strengths of bonds between individuals depending on their age class (adult and juvenile “assortment”). We tested for assortment in edge weights using the assortnet package (version 0.12) (Farine, 2014) to generate an assortment coefficient (*r*, a value from -1 to 1) which we compared to the *r* values of permuted networks. Positive assortment suggests similarly characterised individuals form stronger associations, while negative assortment indicates disassociation (Newman, 2002; Farine, 2014). Following evidence of different levels of associations between the different age groups, we considered if site use patterns uncovered during multistate analysis explained associations between juveniles. In a juvenile-only network we confirmed non-random associations across groups, because of evidence for differential site usage by individuals from our initial multistate analysis which could have structured associations across sites (Farine, 2017).

We also investigated associations within groups to assess if juveniles had non-random associations on a finer scale. We compared the cv values of our network to cv values from permuted networks with data swapped across groups and then within groups. Finally, we explored assortment in association strengths depending on the primary group each juvenile was most commonly recorded in across all surveys, by comparing to the assortment coefficients of permuted networks.

#### Behavioural interactions

To explore how adult and juvenile hihi behaved in groups, for each individual we calculated the proportion of its observations where it was recorded interacting with another bird (separate observations were more than thirty seconds apart) and compared proportions between ages with a Wilcoxon rank sum test. Using proportions accounted for differences in survey effort so that we could combine data from all years. For each juvenile, we then calculated the proportion of total interactions allocated to each behaviour in Table 3.2 and explored if particular types of interactions were correlated using a Principle Components Analysis (PCA) (Budaev, 2010). For any principle components that explained 75% of variance, we next assessed how they correlated with network associations and whether juveniles that behaved in particular ways were more central in the network. We extracted weighted degree scores from our network for each juvenile each year, which explained the number and strength of associations for each bird and thus its placement in the network (animals with more connections tend to be placed more centrally (Krause *et al.*, 2015)). We ranked degrees and divided ranks by the number of juveniles each year, to calculate a proportion rank that was comparable across the different years of the study. We then constructed a GLM with each juvenile’s degree rank as the response and any identified principle components as predictors. To account for non-independence in network data, we generated *P*-values by comparing our observed coefficient to coefficients generated from 1000 models where degree rank values were calculated from permuted networks (Farine and Whitehead, 2015).

**Table 3.** Model-averaged estimates of re-sighting (*ρ*) and movement (*Ψ*) for adult and juvenile hihi in (a) 2015; (b) 2016 and (c) 2017. Estimates generated from multistate models in Supplementary Table 1 which had ΔQAIC weight > 0.00; significant estimates where confidence intervals (LCI, UCI) did not span 0.00 are highlighted in bold. Letters for movement correspond to group sites in Figure 2d, e, f.

| (a) |  | Est. | LCI | UCI |
| --- | --- | --- | --- | --- |
| | $\rho$ Adult | **0.23** | **0.12** | **0.41** |
| | $\rho$ Juvenile | **0.59** | **0.42** | **0.73** |
| | $\psi$ a to b Adult | 0.01 | 0.00 | 0.02 |
| | $\psi$ a to b Juvenile | **0.09** | **0.01** | **0.47** |
| | $\psi$ b to a Adult | 0.00 | -0.01 | 0.02 |
| | $\psi$ b to a Juvenile | 0.02 | 0.00 | 0.15 |
| (b) |  | Est. | LCI | UCI |
| | $\rho$ Adult | **0.17** | **0.09** | **0.31** |
| | $\rho$ Juvenile | **0.36** | **0.22** | **0.53** |
| | $\psi$ b to c Adult | 0.00 | 0.00 | 0.00 |
| | $\psi$ b to c Juvenile | 0.00 | -0.02 | 0.02 |
| | $\psi$ b to d Adult | **0.04** | **0.01** | **0.23** |
| | $\psi$ b to d Juvenile | **0.15** | **0.09** | **0.31** |
| | $\psi$ c to b Adult | 0.00 | -0.01 | 0.02 |
| | $\psi$ c to b Juvenile | 0.02 | 0.00 | 0.11 |
| | $\psi$ c to d Adult | 0.02 | 0.00 | 0.11 |
| | $\psi$ c to d Juvenile | **0.09** | **0.04** | **0.19** |
| | $\psi$ d to c Adult | 0.01 | 0.00 | 0.07 |
| | $\psi$ d to c Juvenile | **0.04** | **0.02** | **0.10** |
| | $\psi$ d to b Adult | 0.01 | -0.01 | 0.03 |
| | $\psi$ d to b Juvenile | **0.04** | **0.01** | **0.23** |
| (c) |  | Est. | LCI | UCI |
| | $\rho$ Adult | **0.22** | **0.17** | **0.28** |
| | $\rho$ Juvenile | **0.44** | **0.40** | **0.49** |
| | $\psi$ b to d Adult | 0.01 | 0.00 | 0.02 |
| | $\psi$ b to d Juvenile | **0.05** | **0.03** | **0.11** |
| | $\psi$ b to e Adult | 0.00 | 0.00 | 0.01 |
| | $\psi$ b to e Juvenile | 0.01 | 0.00 | 0.09 |
| | $\psi$ d to b Adult | 0.00 | 0.00 | 0.01 |
| | $\psi$ d to b Juvenile | **0.02** | **0.01** | **0.04** |
| | $\psi$ d to e Adult | 0.01 | 0.00 | 0.02 |
| | $\psi$ d to e Juvenile | **0.04** | **0.02** | **0.08** |
| | $\psi$ e to b Adult | 0.00 | 0.00 | 0.01 |
| | $\psi$ e to b Juvenile | 0.01 | 0.00 | 0.02 |
| | $\psi$ e to d Adult | **0.04** | **0.01** | **0.13** |
| | $\psi$ e to d Juvenile | **0.20** | **0.11** | **0.33** |

## Results

There were two groups in 2015 and three groups for each of 2016 and 2017, in gully areas (away from feeders) containing water sources and mixed forest. Each year, hihi had multiple associates (mean ± S.E. number of associates: juveniles: 2015 = 15.71 ± 1.59; 2016 = 24.01 ± 1.63; 2017 = 25.29 ± 1.84; adults: 2015 = 8.21 ± 1.00; 2016 = 8.57 ± 1.23; 2017 = 9.84 ± 1.00). The 2015 network represented 379 associations between 33 adults and 31 juveniles; 2016, 1168 associations between 54 adults and 78 juveniles; and 2017, 1400 associations between 61 adults and 87 juveniles. The “gmmevents” event lengths defining associations corresponded to the length of time hihi were re-sighted across consecutive time blocks (median length of event windows (seconds): 2015 = 119.79, 2016 = 90.44, 2017 = 90.75; median re-sighting periods (seconds): 2015 = 90, 2016 = 90, 2017 = 120; Wilcoxon rank sum test comparing length of event windows to re-sighting periods, 2015: *W* = 123240, *P* = 0.06, 2016: *W* = 541210, *P* = 0.54; 2017: *W* = 824380, *P* = 0.17). Both the juvenile/adult and juvenile-only networks showed non-random (preferred and avoided) associations each year (juvenile/adult network: 2015: cv = 2.64, *P*_rand_ = 0.03; 2016: cv = 3.60, *P*_rand_ < 0.001; 2017: cv = 3.56, *P*_rand_ = 0.008; juvenile-only network: 2015: cv = 1.77; 2016: cv = 2.31; 2017: cv = 2.46; in all years, *P*_rand_ values across-location and within-location < 0.001).

### WERE JUVENILES MORE RESIDENT IN GROUPS THAN ADULTS?

There was no difference in the numbers of adults and juveniles detected within and across years (Fisher’s exact test: *N* juveniles = 207; *N* adults = 175; *P* = 0.18). However, juveniles were present on more days than adults (Wilcoxon rank sum test comparing number of days adults and juveniles were re-sighted: 2015: *W* = 235.5, *P* < 0.001; 2015: *W* = 235.5, *P* < 0.001; 2015: *W* = 235.5, *P* < 0.001). Consequently, our multistate analysis estimated that juveniles were re-sighted at least twice as frequently in successive survey occasions compared to adults in all three years (top-ranked models explaining re-sighting included age; Table 3; Figure 2a, b, c; Supplementary Table 1; juveniles *N*: 2015 = 37; 2016 = 79; 2017 = 91; adults *N*: 2015 = 45; 2016 = 62; 2017 = 68). Re-sighting was constant in 2015 and 2017 but varied across survey occasions in 2016 for both adults and juveniles (Supplementary Table 1) suggesting there were small variations in social behavior across years.

**Figure 2.**
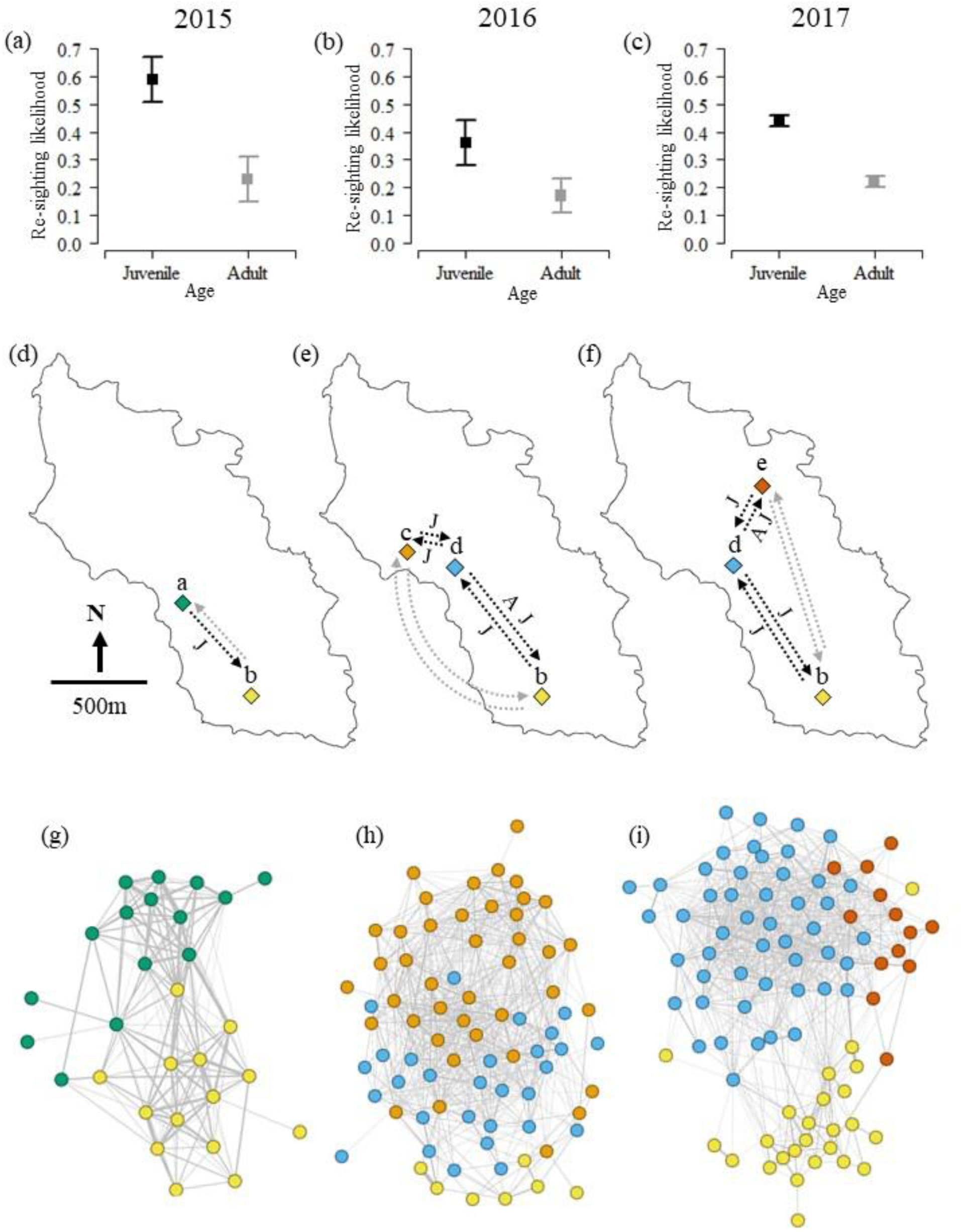
Re-sighting (a-c), movement (d-f) and associations (g-i) for groups (left = 2015; middle = 2016; right = 2017); (a-c) represent mean (± S.E.) re-sighting estimates for juveniles (black lines) and adults (grey lines); (d-f) show movements (dashed lines with arrowheads) between different groups for juveniles (“J”) and adults (“A”) (significant movements from Table 3 are black and lettered, non-significant movements coloured grey); (g-i) show social network diagrams where nodes (circles) represent each hihi and are coloured according to the location in (d-f) where they were seen most often. Lines (edges) represent associations. Strongly associating nodes cluster together more.

Networks reflected these general patterns in residency and showed strong positive assortment by age: each year at least 38% of associations occurred between juveniles only (Table 4; 2015: *r* = 0.15, *P*_rand_ < 0.001; 2016: *r* = 0.25, *P*_rand_ = 0.028; 2017: *r* = 0.19, *P*_rand_ = 0.001). Juveniles were also more likely to interact with others compared to adult hihi (Wilcoxon rank sum test: *W* = 8554.5, *P* < 0.001; although 67/207 juveniles were never observed interacting). Principle Component 1 (PC1) was strongly negatively loaded to “playfight” (Table 5; Supplementary Figure 1), which was the most frequent interaction (mean ± S.E. proportion of total interactions per juvenile that were playfights = 0.25 ± 0.02). Most remaining variation was represented by PC2 and PC3 (Table 5; Supplementary Figure 1). PC2 was loaded most strongly by “huddle” and “chased”, but in opposite directions; this quantified variation in potential affiliative behaviours, because positive scores indicated individuals that huddled more were chased less often. PC3, on the other hand, was loaded negatively by “huddle” and “chased”, but positively by “chase”. This third component described variation where individuals that huddled less chased others more. For individuals that interacted, these three behavioural components did not significantly predict variation in network position (Table 6). However, there was a non-significant tendency that individuals with a more positive PC3 score (more likely to chase, less likely to be chased or huddle) had higher degree ranks (Table 6) suggesting that more dominant individuals may have tended towards being more social.

**Table 4.** Mixing matrices showing distribution of edge weights between adults (“A”) and juvenile (“J”) hihi for (a) 2015, (b) 2016 and (c) 2017 networks. a_i_^w^ are row sums, b_i_^w^ are column sums; due to rounding, sum values may not be exact. Tables are symmetrical so half the values are presented.

| (a) |  |  |  | (b) |  |  |  |
| --- | --- | --- | --- | --- | --- | --- | --- |
| | A | J | $a_i^w$ | | A | J | $a_i^w$ |
| **A** | 0.208 | - | 0.414 | **A** | 0.121 | - | 0.268 |
| **J** | 0.206 | 0.381 | 0.587 | **J** | 0.146 | 0.586 | 0.733 |
| **$b_i^w$** | 0.414 | 0.587 | 1.000 | **$b_i^w$** | 0.268 | 0.733 | 1.000 |

| (c) |  |  |  |
| --- | --- | --- | --- |
| | A | J | $a_i^w$ |
| **A** | 0.111 | - | 0.273 |
| **J** | 0.162 | 0.565 | 0.727 |
| **$b_i^w$** | 0.273 | 0.727 | 1.000 |

**Table 5.**
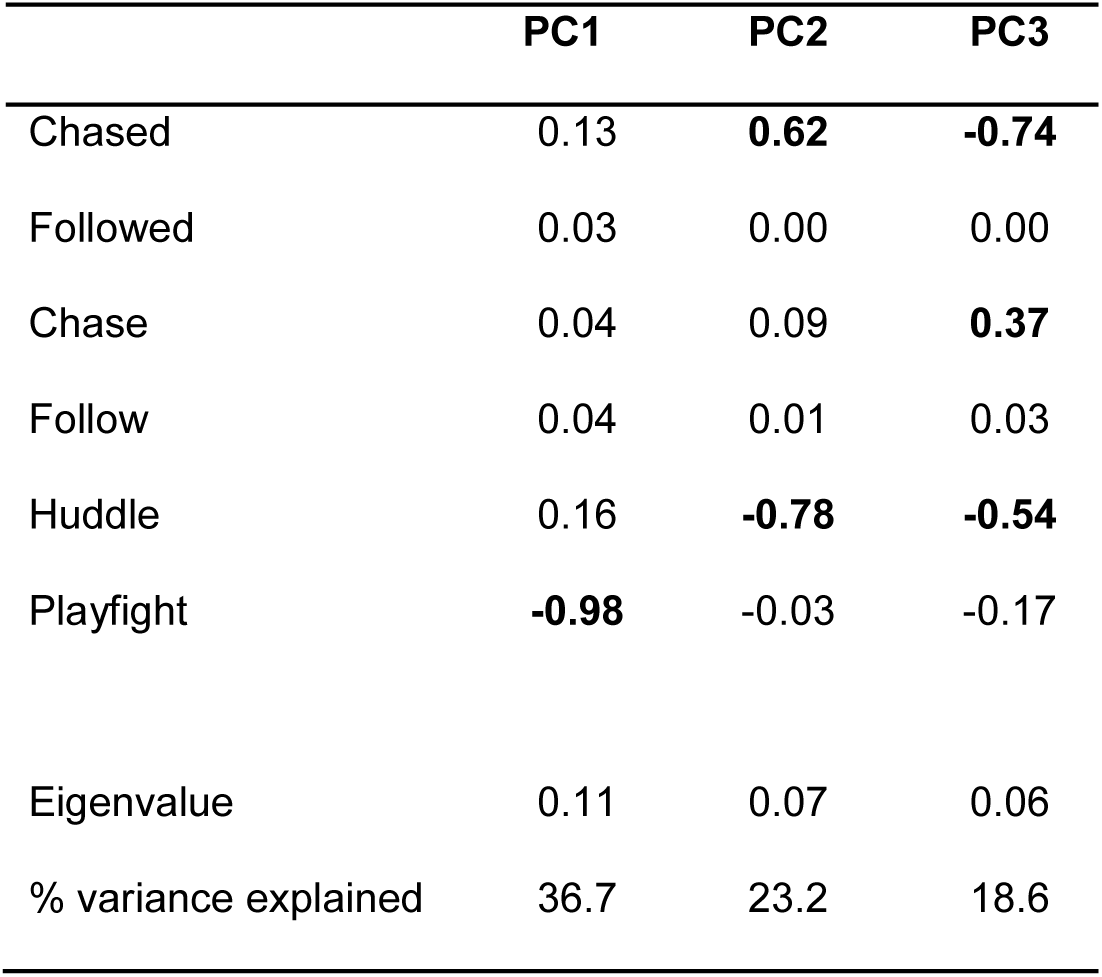
Principle components analysis (PCA) of juvenile social behaviours seen in group sites. The first three components accounted for more than 75% of variance (components 4-6 accounted for 21.5% variance in total and are not presented). Behaviours that loaded most on each PC are highlighted in bold.

**Table 6.** Results of a binomial GLM analysing variation in degree rank depending on PC1, PC2, and PC3 describing variation in interactions between juvenile hihi (Table 5). Coefficients, standard errors and *z* values are presented. Both the *P-*value of the model and the *P-*value generated using coefficients from 1000 randomised networks (specified as *P*_rand_) are presented, for comparison. Marginal significance of PC3 indicated with “.”.

|  |  | coeff. | S.E. | z-value | *P*-value | *P*<sub>rand</sub> |
| --- | --- | --- | --- | --- | --- | --- |
| degree ~ | intercept | 0.82 | 0.21 | 3.92 | < 0.001 | 1.00 |
|  | PC1 | -0.53 | 0.55 | -0.97 | 0.33 | 0.87 |
|  | PC2 | 0.33 | 0.57 | 0.57 | 0.57 | 0.49 |
|  | PC3 | 0.56 | 0.81 | 0.68 | 0.50 | 0.06 . |

### DID GROUPS FORM IN STABLE LOCATIONS, OR DID THEY MOVE?

Quantifying movement (*Ψ*) in our multistate analysis showed a low likelihood that hihi transitioned between group sites, although this did vary depending on where birds were moving to and from (Table 3, Figure 2d, e, f; Supplementary Table 1). Movement also depended on age, and some juveniles did move groups between each survey (Table 3; Figure 2d, e, f; Supplementary Table 1). However, on average only two or three juveniles moved between each survey (mean: 2015 = 2; 2016 = 3; 2017 = 3), and movement also varied among individuals (maximum number of moves per individual: 2015 = 3; 2016 = 7; 2017 = 7; juveniles that never moved groups: 2015 = 29/37; 2016 = 35/79; 2017 = 56/91). Furthermore, in the social network analysis we found that juvenile-only networks showed strong positive assortment by primary group in all three years, while associations among juveniles resident in different sites were much weaker (Table 7; Figure 2g, h, i; 2015: *r* = 0.513, *P*_rand_ < 0.001; 2016: *r* = 0.32, *P*_rand_ < 0.001; 2017: *r* = 0.58, *P*_rand_ < 0.001).

**Table 7.** Mixing matrices showing distribution of edge weights between juveniles depending on the group where they were most commonly located (site lettering refers to group locations in Figure 6) in (a) 2015, (b) 2016 and (c) 2017. ai^w^ are the row sums, bi^w^ are the column sums; due to rounding, sum values may not be exact. Tables are symmetrical so half the values are presented.

| (a) | Site a | Site b | $a_i^w$ | (b) | Site c | Site d | Site b | $a_i^w$ |
| --- | --- | --- | --- | --- | --- | --- | --- | --- |
| Site a | 0.330 | - | 0.450 | Site c | 0.503 | - | - | 0.653 |
| Site b | 0.121 | 0.429 | 0.550 | Site d | 0.141 | 0.144 | - | 0.298 |
| $b_i^w$ | 0.450 | 0.550 | 1.000 | Site b | 0.009 | 0.013 | 0.027 | 0.049 |
| | | | | $b_i^w$ | 0.653 | 0.298 | 0.049 | 1.000 |

| (c) | Site d | Site b | Site e | $a_i^w$ |
| --- | --- | --- | --- | --- |
| Site d | 0.575 | - | - | 0.674 |
| Site b | 0.028 | 0.155 | - | 0.189 |
| Site e | 0.071 | 0.006 | 0.061 | 0.137 |
| $b_i^w$ | 0.674 | 0.189 | 0.137 | 1.000 |

### WERE JUVENILES RELATED TO ADULTS AND OTHER JUVENILES?

In the re-sighting data each year there were very few occasions when juveniles were seen in the same group during the same survey as their parents (mean ± S.E. proportion of surveys: 2015 = 0.02 ± 0.02; 2016 = 0.03 ± 0.01; 2017 = 0.08 ± 0.02), or their maternal-siblings (Figure 3a; mean ± S.E. proportion of surveys: 2015 = 0.22 ± 0.08; 2016 = 0.25 ± 0.04; 2017 = 0.28 ± 0.04). Individuals that grouped closer to their nest-of-origin were not more likely to be seen with maternal-siblings each year (Figure 3b; null model highest ranked; Supplementary Table 2). Being recorded in more surveys also did not affect cooccurrence with maternal-siblings in any year (Supplementary Table 2). Together, this low likelihood of juveniles being resident with parents or maternal-siblings suggested that these individuals had very limited opportunities to associate, so we did not analyse assortment by relatedness in networks.

**Figure 3.**
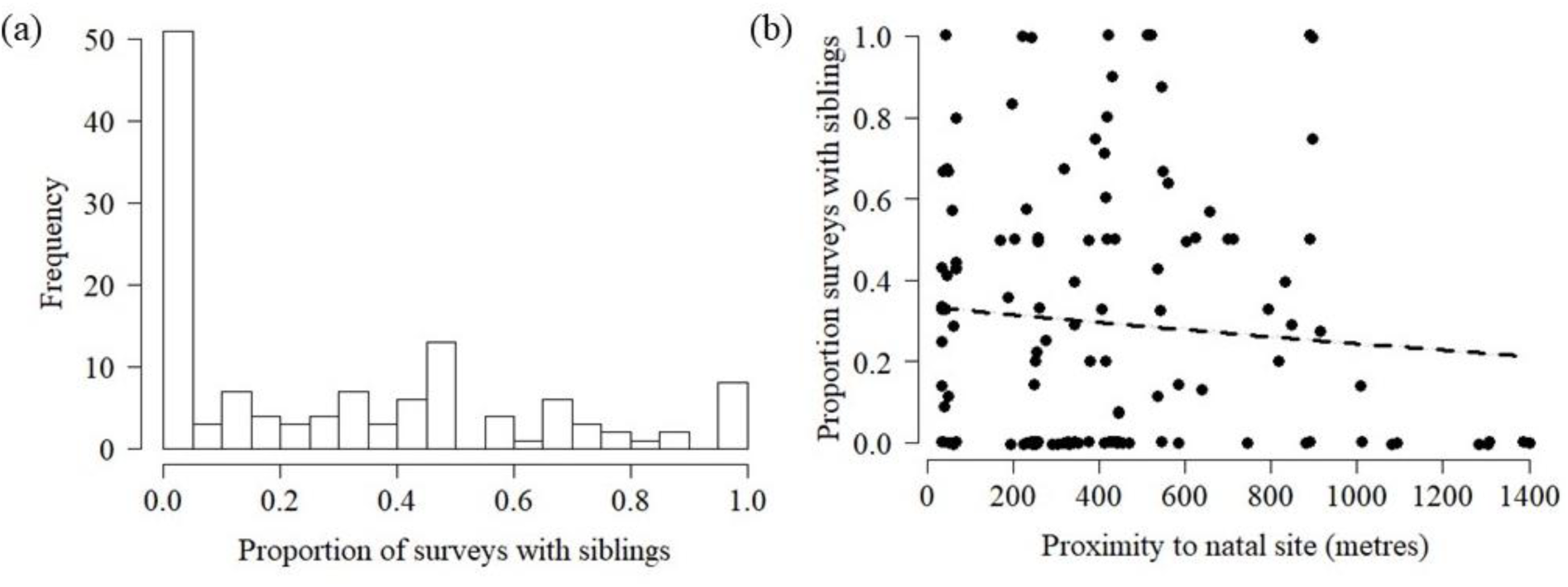
(a) Variation among juveniles in the proportion of occasions where they were recorded with their maternal-siblings (all years of the study included), and (b) relationship between proportion of occasions seen with siblings and the distance from each juvenile’s natal site to where they grouped. In (b), confidence intervals were too narrow to plot (Supplementary Table 2).

## Discussion

Here we used both a multistate analysis and social network analysis approach to characterise the location use, age composition, and relatedness of hihi groups that form at the end of each breeding season. We found that groups formed non-randomly and occurred in consistent locations within each year, with little movement across our study site. Multistate analysis indicated that groups were formed predominantly of juveniles, and although some adults were observed their presence was more transient. Network associations reflected these differences in residency: rather than associating with adults, juveniles most strongly associated with other juveniles frequently present in the same group locations. Juveniles also interacted more frequently with other birds compared to adults. However, despite differences among individuals in the amount of affiliative- or aggressive-type interactions, the types of behavioural interactions did not significantly predict a juveniles’ number of network associates. Finally, juveniles were almost never seen with their parents (occurred in only 2-8% of surveys across the study) and were also re-sighted without their nest mates in the majority (72-78%) of surveys. Together, these results suggest juvenile hihi groups most closely resemble the “gangs” described in juvenile ravens by Dall and Wright (2009) (Table 3.1), where juvenile birds aggregate around communal roosts (Wright, Stone and Brown, 2003) or other social meeting places (Ward and Zahavi, 2008) which are separate from main colonies and thus have limited interaction with adults. This is in contrast to flocks, which can move over large distances (Templeton *et al.*, 2012) or crèches, where juveniles associate with related adults (Balda and Balda, 1978).

Animals can aggregate if ecological factors (such as rich foraging grounds) cause them to coexist in the same place at the same time (Mourier, Vercelloni and Planes, 2012; Strandburg-Peshkin *et al.*, 2017; Gall and Manser, 2018), but location use can also arise as a consequence of preferring to associate with others (Fletcher, 2007; Firth and Sheldon, 2016). The aggregations of juvenile hihi we detected here could have been a by-product of differential habitat use according to age to avoid competition with more dominant adults (Catterall, Kikkawa and Gray, 1989; Marchetti and Price, 1989; Sol *et al.*, 1998). Alternatively, groups could have arisen through juveniles choosing to associate with individuals of a similar phenotype (Croft *et al.*, 2005). Understanding the intricate link between current environment and how or why associations form is still a fledgling topic in social network analysis (Madden *et al.*, 2009; Godfrey, Sih and Bull, 2013; Pinter-Wollman *et al.*, 2013; Leu *et al.*, 2016). We did not explicitly test this link here (for example, by changing the environment and comparing network structures (Formica *et al.*, 2016)) so cannot fully conclude if ecology or individual choice determined group associations. However, repeating observations across years did show similar characteristics in groups and their locations, albeit with a small level of variation, perhaps suggesting climatic conditions or other ecological variables of the sites affected group formation (Krause and Ruxton, 2002). Thus, the value of long term studies is that they allow for replicates that demonstrate whether the same determinants structure animal groups across years (Shizuka *et al.*, 2014) especially when individual identities differ year-on-year (as in our study, with different juvenile cohorts).

Regardless of whether groups arose due to active choice by individuals or a more incidental aggregation based on environment, the result was that non-random associations formed between juveniles which could mediate behaviours such finding food, and avoiding predators or disease (Krause and Ruxton, 2002; Krause, Lusseau and James, 2009; Drewe, 2010; Aplin *et al.*, 2012; Schakner *et al.*, 2017). As we found limited co-occurrence and associations between differently-aged animals, are there benefits to aggregating with other juveniles rather than adults? This could be a potentially risky strategy as young animals are naïve (Galef and Laland, 2005) and do not always behave appropriately to suit the current environment (Clayton, 1994). Individuals appear to recognise these risks in some species (such as capuchin monkeys, *Cebus apella*) and prefer to pay attention to adults rather than juveniles if given the choice (Ottoni, De Resende and Izar, 2005). However, in gang-type groups the limited presence of adults creates little opportunity to associate with these more experienced individuals, in contrast to flocks (Templeton *et al.*, 2012) or crèches (Heinsohn, 1991).

While young animals by themselves may be naïve, large groups of juveniles are still thought to be beneficial because they can act as “information centres” (Dall and Wright, 2009) where associating with many animals collectively gathering and sharing information may help overcome any one individual’s inexperience (Ward and Zahavi, 2008). For example, in quelea (*Quelea quelea*), parents leave their young after approximately three weeks of care, and young then form assemblages which help them to exploit their habitat and forage successfully without learning from adults (Ward and Zahavi, 2008). Similarly, juvenile raven gangs respond collectively to new, ephemeral, food sources (Marzluff, Heinrich and Marzluff, 1996; Dall and Wright, 2009). Furthermore, in this context other factors such as relatedness may not be important for grouping because non-kin provide a broader range of information collected from different experiences, which could be more relevant to the current environment (Schwab, Bugnyar and Kotrschal, 2008; Kulahci *et al.*, 2016). Young animals are known to pay more attention to non-kin particularly when early life conditions were suboptimal, suggesting they adjust associations depending on payoff (Farine, Spencer and Boogert, 2015). Hihi do have high rates of extra-pair paternity (Brekke *et al.*, 2013), and unfortunately genetic data was not available at the time of the study, but the general low presence of adults or half-siblings suggests relatedness was not important to their grouping (Saitou, 1978, 1979; Hirsch *et al.*, 2013; Arnberg *et al.*, 2015). Overall, if hihi juvenile groups may be information centres then it will be valuable to test how they inform foraging behaviour.

While aggregating, juvenile hihi interacted directly with other individuals. Some behaviours were not equal (for example, a hihi that was chased did not then become the chaser) and so could be establishing dominance in these groups (Drews, 1993). Other behaviours appeared to be affiliative, and consistent with definitions of social play (Diamond and Bond, 2003). Social play is known in other gang-forming juveniles (ravens) (Heinrich and Smolker, 1998; Diamond and Bond, 2003) and is generally thought to be a more complex behaviour associated with large brain sizes, but previous reviews have cautioned that its apparent absence in other species could be due to a lack of research (Diamond and Bond, 2003). Interactions between juveniles have been suggested to be one route by which information is shared in other species (Diamond and Bond, 2003). However, we did not find a link between likelihood of interacting and network position (degree). This may indicate interactions and familiarity between specific individuals are not crucial to information dissemination in young hihi (Schwab, Bugnyar and Kotrschal, 2008; Guillette, Scott and Healy, 2016; Ramakers *et al.*, 2016), as an individual’s number of associates can be important for information acquisition (Aplin *et al.*, 2012; Snijders *et al.*, 2014). As yet, it remains unclear what structures juvenile hihi network position in groups, so further work is needed to test why groups form and how this influences sociality, to help further understand the importance of group structure for learning in young birds.

To conclude, we show that juvenile hihi are commonly found in groups during their first few months of independence from parents. These groups form in spatially-separated locations and are dominated by juveniles, with little opportunity to interact with adults. The structure of gang-like groups in young hihi create the potential for many naïve individuals to associate, and potentially share information, Next, it will be valuable to test more explicitly whether these groups inform behaviour in young hihi. By doing so, we can explore if such groups provide opportunities to help young birds overcome any one individual’s disadvantage of being naïve, or whether there are downsides of associating with inexperienced peers.

## Acknowledgements

We are grateful to the Department of Conservation and Supporters of Tiritiri Matangi for their permission to conduct this research.

